# Evaluating tree biomass estimation in trans-Atlantic mangrove species: comparing bole diameter measurements for improved accuracy

**DOI:** 10.1101/2025.04.19.649682

**Authors:** Charles A. Price, Todd A. Schroeder, Benjamin Branoff, Humfredo Mercano-Vega, Nicole Pilot-Torres, Morgan Chaudry, Michael Ross, Monica Papeş, Skip Van Bloem

**Author notes:** **Corresponding Author:** Charles A. Price, National Institute for Modelling Biological Systems, Ecology and Evolutionary Biology, University of Tennessee, Knoxville, TN, 37996-3140, USA.

## Abstract

Estimating the biomass of terrestrial forests generally, and mangrove forests in particular, is an area of considerable interest. Most approaches rely on empirically derived allometric models to predict tree biomass. The single parameter with the strongest predictive ability in most studies is diameter at breast height (DBH), however the use of DBH arose primarily out of convenience, not from an analysis of tree form. While DBH explains a lot of variability in other tree metrics such as height or above ground biomass, its utility in smaller species is uncertain. Here we used measurements from 302 destructively sampled mangrove trees of four species to test which of three bole diameter measurements, basal stem diameter (BSD), diameter at 30 cm (D30), and DBH, is the best predictor of above ground biomass. D30 had the highest mean coefficient of determination (*R*^*2*^) and lowest mean root mean squared error (RMSE) across all site/species combinations. However, the improvement over DBH was modest, with a mean across all site/species combinations of 1.58 kg RMSE and *R*^*2*^ of 0.948 for D30, compared to 1.63 kg RMSE and *R*^*2*^ of 0.917 for DBH. Nevertheless, D30 may have utility in future studies as it allows for lower size thresholds and has better overall explanatory power than DBH.

## Introduction

Accurate prediction of the aboveground biomass of terrestrial trees underpins research at numerous levels, from interspecific resource exchange in tree communities to global models of forest dynamics [1]. Foresters and plant ecologists have long used diameter at breast height (DBH) to estimate biomass. However, the use of DBH as a standard measurement point emerged from a combination of convenience and sociocultural factors [2], rather than from an objective analysis of tree form. Moreover, the height at which DBH is measured differs across studies [3].

Despite these historical idiosyncrasies, DBH has been shown to correlate well with tree height and aboveground mass for large trees, particularly those with large boles growing in closed canopy forests [1]. However, for smaller woody trees and shrubs, it may not be the best predictor. Many small trees and shrubs branch extensively below DBH height, thus DBH for those plants captures the biomechanical and hydraulic investment for some, but not all of the plant. Foresters working in arid environments have long understood this and have measured the diameter at a woody plants’ base instead of DBH as a correlate of above ground biomass. For example, the US Nationwide Forest Inventory (NFI) program [4] measures the diameter at root collar (DRC) for woodland species (e.g., pinyon-juniper, mesquite, oak woodlands, etc.) growing in arid regions of the western U.S. [5].

Mangrove species growing in the US mainland and Caribbean islands exhibit a wide range of growth patterns, ranging from tall trees with large and easily measured boles, to small shrubbier forms, sometimes referred to as dwarf mangroves [6]. Mangrove growth and tree size are known to vary at geographic scales [7, 8], and can be influenced by surface hydrology and soil nutrient composition [9-11], temperature, humidity and salinity [12, 13], and disturbance frequency and intensity [14, 15]. Toward the northern extent of mangroves’ range, their growth is strongly influenced by cold temperatures, with shorter growth seasons leading to mangroves that tend to be smaller and shrubbier [6, 16].

As with other terrestrial forest systems, empirical allometric equations used to derive biomass have played a central role in the estimation of mangrove biomass from community to landscape scales [8]. Numerous studies worldwide have harvested mangrove trees and generated locally derived allometric functions [6, 7, 11, 17-24]. While some studies have, out of necessity, included diameter measurement points lower than DBH [6, 19], we are not aware of any attempts to systematically evaluate which of several possible measurement points has the greatest predictive power.

Here we utilize data from two sources that allows us to evaluate which of three measurement points at the basal end of trees is the best predictor of above ground biomass. The first dataset (San Juan dataset) includes recent measurements for the linear dimensions (height and canopy diameter) and aboveground mass of 152 individual trees collected in San Juan, Puerto Rico (Price et al. in prep). The second dataset (Biscayne dataset) comes from an analysis of mangrove form and allometry using 150 individuals collected in Biscayne, Florida, USA [6]. Both studies contain data for the three true mangrove species growing regionally, *Avicennia germinans* (L.)L., *Laguncularia racemosa* (L.)C.F.Gaertn., and *Rhizophora mangle* L.; in addition, the San Juan dataset includes a common mangrove associate, *Conocarpus erectus* L. Tree diameter was measured at the base of the stem (BSD), at 30 cm above base height (D30), and at breast height (DBH) for both datasets, which we further analyze to determine the best predictor of aboveground mass.

## Materials and Methods

The San Juan dataset was collected in November 2024 (see Price et al. in prep for full methods). Collection methods for the Biscayne dataset can be found in Ross et al. [6]. As both studies collect similar measurements using similar methods, we only briefly summarize the protocol used to collect the San Juan dataset.

For each tree, diameter was measured at a point just above the butt swell (*BSD*), at 30 cm (*D30*) above the *BSD* measurements point, and at 130 cm above the *BSD* measurement point (referred to here as *DBH*). The height of each tree was measured with a tape measure (short trees) or a hypsometer (tall trees). The canopy diameter was measured at its widest point, and again at an axis perpendicular to the first canopy measurement. All trees were cut at ground level and the wet mass of each tree was measured with either a digital scale (small trees) or a spring scale (large trees) suspended from a tall aluminum step-ladder. For large trees, each individual was cut into several sections to enable weighing.

### Wood Discs

Due to logistical constraints, we were unable to dry and weigh entire trees to determine dry biomass. Instead, we estimated the dry mass fraction from wood discs cut from branches of various sizes for each species; the discs were dried to constant mass and weighed to calculate the dry mas fraction as dry mass/wet mass. We then multiplied *wet mass* of the tree by the dry mass fraction of the wood discs to yield aboveground *dry mass* estimates. This approach is quite similar to that used by several other researchers reporting biomass allometries for these species [11, 20-22, 24].

### Allometric Analyses

Bivariate relationships between all measurements were estimated using ordinary least squares regression (OLS). To estimate regression function variables and model fits we utilized the software package Standardized Major Axis Tests and Routines (SMATR) [25, 26]. OLS regression is typically used in allometric studies where one wants to predict the values of the Y-variable using the X-variable [26]. Data were log transformed prior to regression fitting to meet the assumption of homogeneity of variance. All regression functions were tested at the p< 0.05 significance level.

We fit OLS regression functions to multiple different groupings, both inter and intraspecifically, within and between sites. To examine patterns across both the Biscayne and San Juan data, we combined both datasets (referred to as Combined). We also analyzed intraspecific data both within and between sites (Table 1). Not all trees have all three measurements as some trees were simply too small to measure DBH or even D30. To determine if our findings were influenced by samples with only one or two of the measurements (BSD or D30), we also generated regression functions using only trees with all three measurements (denoted with a “3” in Table 2).

**Table 1.**
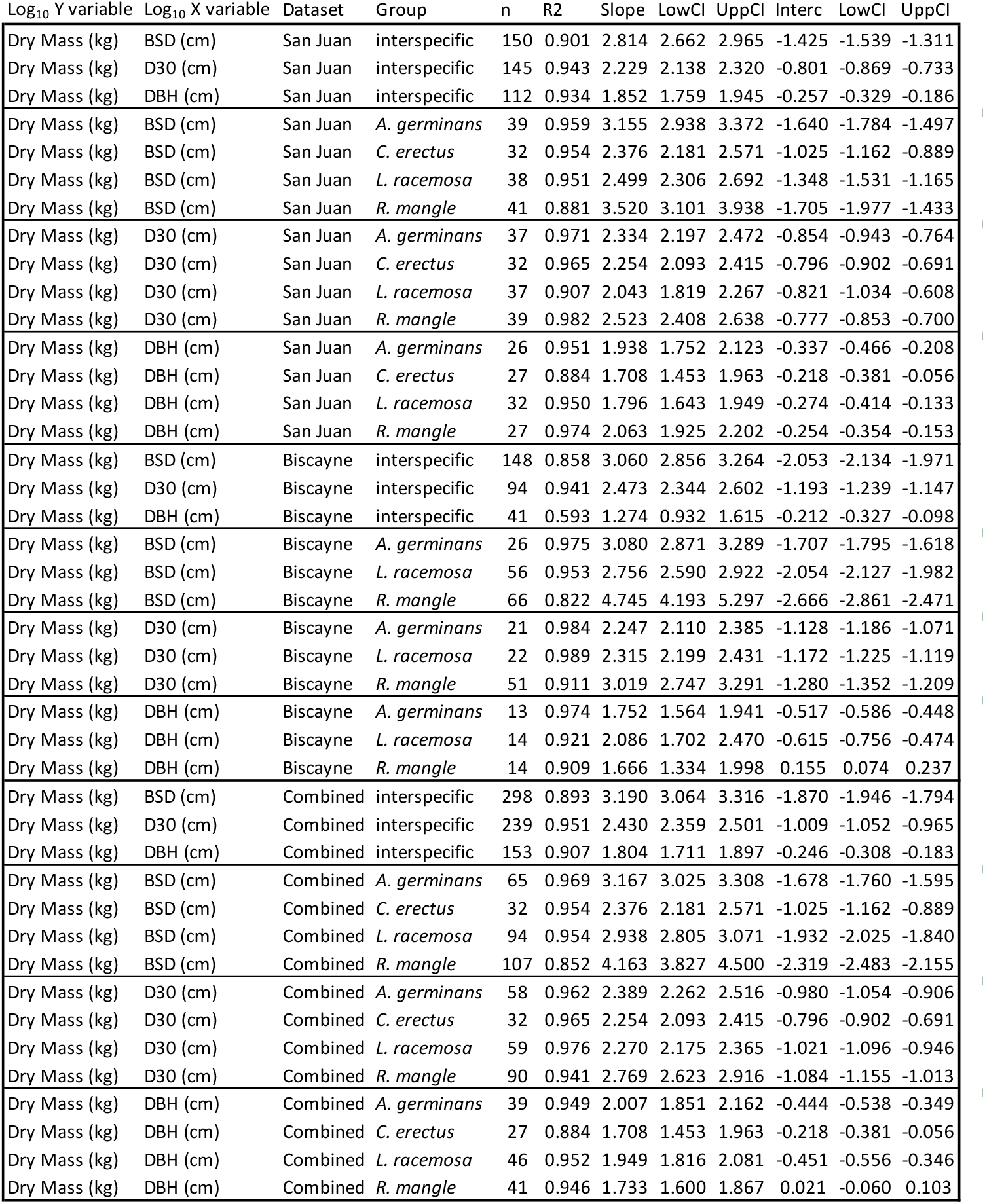
OLS allometric regression data. The first and second columns contain the Log10 transformed Y and X variables, respectively, followed by the dataset, group, sample size (n), R2 values, slope with lower and upper 95% confidence intervals, and intercept with lower and upper 95% confidence intervals. Thick borders are included to help the reader identify data analyzed at the same level.

| Log<sub>10</sub> Y variable | Log<sub>10</sub> X variable | Dataset | Group | n | R<sup>2</sup> | Slope | LowCI | UppCI | Interc | LowCI | UppCI |
| --- | --- | --- | --- | --- | --- | --- | --- | --- | --- | --- | --- |
| Dry Mass (kg) | BSD (cm) | San Juan | interspecific | 150 | 0.901 | 2.814 | 2.662 | 2.965 | -1.425 | -1.539 | -1.311 |
| Dry Mass (kg) | D30 (cm) | San Juan | interspecific | 145 | 0.943 | 2.229 | 2.138 | 2.320 | -0.801 | -0.869 | -0.733 |
| Dry Mass (kg) | DBH (cm) | San Juan | interspecific | 112 | 0.934 | 1.852 | 1.759 | 1.945 | -0.257 | -0.329 | -0.186 |
| Dry Mass (kg) | BSD (cm) | San Juan | *A. germinans* | 39 | 0.959 | 3.155 | 2.938 | 3.372 | -1.640 | -1.784 | -1.497 |
| Dry Mass (kg) | BSD (cm) | San Juan | *C. erectus* | 32 | 0.954 | 2.376 | 2.181 | 2.571 | -1.025 | -1.162 | -0.889 |
| Dry Mass (kg) | BSD (cm) | San Juan | *L. racemosa* | 38 | 0.951 | 2.499 | 2.306 | 2.692 | -1.348 | -1.531 | -1.165 |
| Dry Mass (kg) | BSD (cm) | San Juan | *R. mangle* | 41 | 0.881 | 3.520 | 3.101 | 3.938 | -1.705 | -1.977 | -1.433 |
| Dry Mass (kg) | D30 (cm) | San Juan | *A. germinans* | 37 | 0.971 | 2.334 | 2.197 | 2.472 | -0.854 | -0.943 | -0.764 |
| Dry Mass (kg) | D30 (cm) | San Juan | *C. erectus* | 32 | 0.965 | 2.254 | 2.093 | 2.415 | -0.796 | -0.902 | -0.691 |
| Dry Mass (kg) | D30 (cm) | San Juan | *L. racemosa* | 37 | 0.907 | 2.043 | 1.819 | 2.267 | -0.821 | -1.034 | -0.608 |
| Dry Mass (kg) | D30 (cm) | San Juan | *R. mangle* | 39 | 0.982 | 2.523 | 2.408 | 2.638 | -0.777 | -0.853 | -0.700 |
| Dry Mass (kg) | DBH (cm) | San Juan | *A. germinans* | 26 | 0.951 | 1.938 | 1.752 | 2.123 | -0.337 | -0.466 | -0.208 |
| Dry Mass (kg) | DBH (cm) | San Juan | *C. erectus* | 27 | 0.884 | 1.708 | 1.453 | 1.963 | -0.218 | -0.381 | -0.056 |
| Dry Mass (kg) | DBH (cm) | San Juan | *L. racemosa* | 32 | 0.950 | 1.796 | 1.643 | 1.949 | -0.274 | -0.414 | -0.133 |
| Dry Mass (kg) | DBH (cm) | San Juan | *R. mangle* | 27 | 0.974 | 2.063 | 1.925 | 2.202 | -0.254 | -0.354 | -0.153 |
| Dry Mass (kg) | BSD (cm) | Biscayne | interspecific | 148 | 0.858 | 3.060 | 2.856 | 3.264 | -2.053 | -2.134 | -1.971 |
| Dry Mass (kg) | D30 (cm) | Biscayne | interspecific | 94 | 0.941 | 2.473 | 2.344 | 2.602 | -1.193 | -1.239 | -1.147 |
| Dry Mass (kg) | DBH (cm) | Biscayne | interspecific | 41 | 0.593 | 1.274 | 0.932 | 1.615 | -0.212 | -0.327 | -0.098 |
| Dry Mass (kg) | BSD (cm) | Biscayne | *A. germinans* | 26 | 0.975 | 3.080 | 2.871 | 3.289 | -1.707 | -1.795 | -1.618 |
| Dry Mass (kg) | BSD (cm) | Biscayne | *L. racemosa* | 56 | 0.953 | 2.756 | 2.590 | 2.922 | -2.054 | -2.127 | -1.982 |
| Dry Mass (kg) | BSD (cm) | Biscayne | *R. mangle* | 66 | 0.822 | 4.745 | 4.193 | 5.297 | -2.666 | -2.861 | -2.471 |
| Dry Mass (kg) | D30 (cm) | Biscayne | *A. germinans* | 21 | 0.984 | 2.247 | 2.110 | 2.385 | -1.128 | -1.186 | -1.071 |
| Dry Mass (kg) | D30 (cm) | Biscayne | *L. racemosa* | 22 | 0.989 | 2.315 | 2.199 | 2.431 | -1.172 | -1.225 | -1.119 |
| Dry Mass (kg) | D30 (cm) | Biscayne | *R. mangle* | 51 | 0.911 | 3.019 | 2.747 | 3.291 | -1.280 | -1.352 | -1.209 |
| Dry Mass (kg) | DBH (cm) | Biscayne | *A. germinans* | 13 | 0.974 | 1.752 | 1.564 | 1.941 | -0.517 | -0.586 | -0.448 |
| Dry Mass (kg) | DBH (cm) | Biscayne | *L. racemosa* | 14 | 0.921 | 2.086 | 1.702 | 2.470 | -0.615 | -0.756 | -0.474 |
| Dry Mass (kg) | DBH (cm) | Biscayne | *R. mangle* | 14 | 0.909 | 1.666 | 1.334 | 1.998 | 0.155 | 0.074 | 0.237 |
| Dry Mass (kg) | BSD (cm) | Combined | interspecific | 298 | 0.893 | 3.190 | 3.064 | 3.316 | -1.870 | -1.946 | -1.794 |
| Dry Mass (kg) | D30 (cm) | Combined | interspecific | 239 | 0.951 | 2.430 | 2.359 | 2.501 | -1.009 | -1.052 | -0.965 |
| Dry Mass (kg) | DBH (cm) | Combined | interspecific | 153 | 0.907 | 1.804 | 1.711 | 1.897 | -0.246 | -0.308 | -0.183 |
| Dry Mass (kg) | BSD (cm) | Combined | *A. germinans* | 65 | 0.969 | 3.167 | 3.025 | 3.308 | -1.678 | -1.760 | -1.595 |
| Dry Mass (kg) | BSD (cm) | Combined | *C. erectus* | 32 | 0.954 | 2.376 | 2.181 | 2.571 | -1.025 | -1.162 | -0.889 |
| Dry Mass (kg) | BSD (cm) | Combined | *L. racemosa* | 94 | 0.954 | 2.938 | 2.805 | 3.071 | -1.932 | -2.025 | -1.840 |
| Dry Mass (kg) | BSD (cm) | Combined | *R. mangle* | 107 | 0.852 | 4.163 | 3.827 | 4.500 | -2.319 | -2.483 | -2.155 |
| Dry Mass (kg) | D30 (cm) | Combined | *A. germinans* | 58 | 0.962 | 2.389 | 2.262 | 2.516 | -0.980 | -1.054 | -0.906 |
| Dry Mass (kg) | D30 (cm) | Combined | *C. erectus* | 32 | 0.965 | 2.254 | 2.093 | 2.415 | -0.796 | -0.902 | -0.691 |
| Dry Mass (kg) | D30 (cm) | Combined | *L. racemosa* | 59 | 0.976 | 2.270 | 2.175 | 2.365 | -1.021 | -1.096 | -0.946 |
| Dry Mass (kg) | D30 (cm) | Combined | *R. mangle* | 90 | 0.941 | 2.769 | 2.623 | 2.916 | -1.084 | -1.155 | -1.013 |
| Dry Mass (kg) | DBH (cm) | Combined | *A. germinans* | 39 | 0.949 | 2.007 | 1.851 | 2.162 | -0.444 | -0.538 | -0.349 |
| Dry Mass (kg) | DBH (cm) | Combined | *C. erectus* | 27 | 0.884 | 1.708 | 1.453 | 1.963 | -0.218 | -0.381 | -0.056 |
| Dry Mass (kg) | DBH (cm) | Combined | *L. racemosa* | 46 | 0.952 | 1.949 | 1.816 | 2.081 | -0.451 | -0.556 | -0.346 |
| Dry Mass (kg) | DBH (cm) | Combined | *R. mangle* | 41 | 0.946 | 1.733 | 1.600 | 1.867 | 0.021 | -0.060 | 0.103 |

**Table 2.**
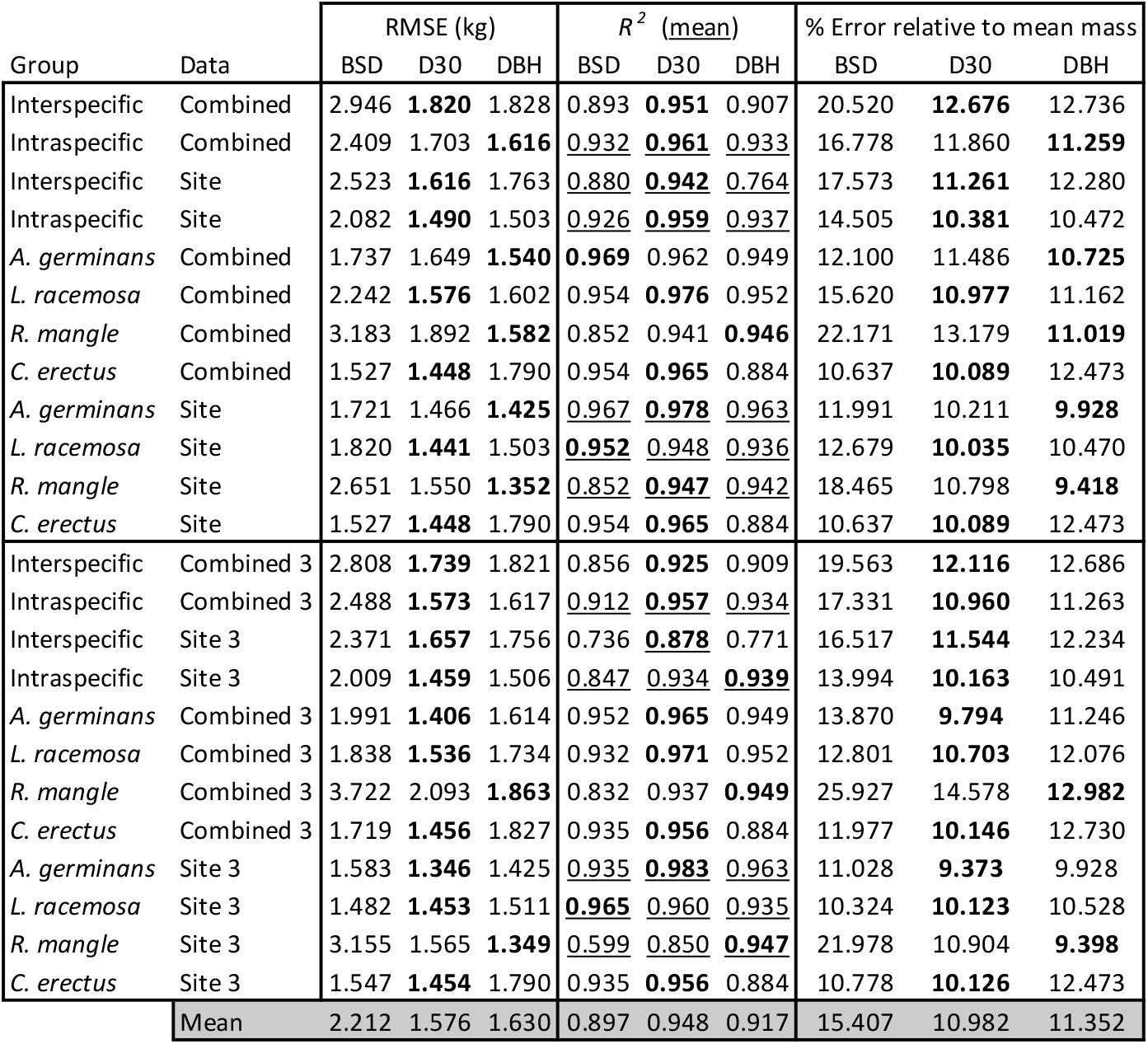
*RMSE* and *R*^*2*^ for different data groupings. The first column contains the species composition (inter, intra, or species specific), and the second column the data composition for each statistic reported in the remaining columns. Columns 3-5 contain the RMSE (kg), columns 6-8 the *R*^*2*^, and columns 9-11 the % error relative to the mean biomass in the dataset for BSD, D30 and DBH, respectively (see Methods). The lowest value in a row for each metric is in bold. Values that represent means are underlined. The mean for each column is given in the final row. Thick borders are included to help the reader identify data analyzed at the same level.

### Root Mean Squared Error

To determine which measure of tree diameter (BSD, D30, DBH) was the best predictor of total aboveground biomass (wet mass or dry mass), and to compare results between regression models at different grouping levels (interspecifically or intraspecifically) we calculated the root mean squared error (RMSE) using standard formula, which represents the mean difference between the values predicted by the regression model and the actual measured values.

#### Diameter Ratios

To examine whether there was a systematic change in the values of our three diameter measurements, relative to one another, we looked at how their ratios (BSD/DBH, BSD/D30, D30/DBH) changed as a function of tree height. We also plotted the diameter measurements against one another to see if regression slopes fit to those data (standardized major axis regression) differed from the simple null expectation that they have a slope of 1 (isometric change).

## Results

We were able to collect 41 *R. mangle*, 40 *A. germinans*, 39 *L. racemosa*, and 32 *C. erectus* individual trees from the site in San Juan. The data from Ross et al. [6] contained measurements for 66 *R. mangle*, 26 *A. germinans*, and 56 *L. racemosa* individuals.

Allometric regression results for the relationships between wet or dry mass, and our three different diameter measurements (BSD, D30, and DBH) are shown in Fig. 1 and Table 1. Overall, there is very little variability around the fitted linear relationships between tree diameter and mass measurements. The mean *R*^*2*^ across all the relationships in Table 1 is 0.928. *R*^*2*^ values for different data groupings are provided in Table 2. Among the three diameter measurements, D30 had the highest *R*^*2*^ value in 17/24 cases and DBH had the highest *R*^*2*^ value in 7/24 cases. All three diameter ratios decreased with increasing tree size (Fig 2) and BSD, D30, and DBH were all strongly correlated with one another (Fig 3).

**Figure 1.**
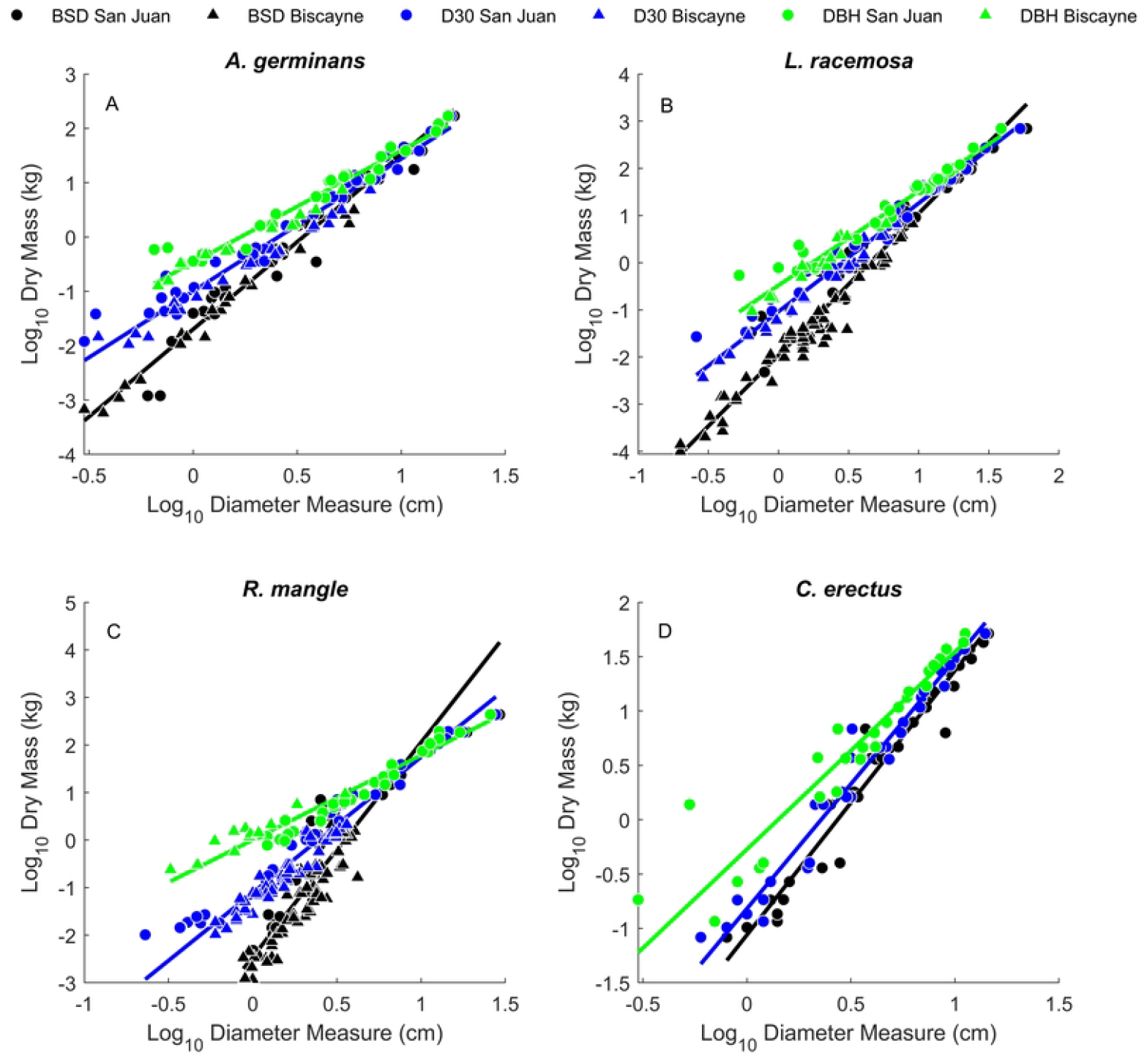
Bivariate relationships between the three diameter measurements (BSD, D30, and DBH) and above ground dry mass for the four mangrove species. Each panel corresponds to data for a given species as denoted in the panel title. Symbols with a red perimeter are those individual trees for which all three measurements were collected. Notice that the three functions appear to converge for larger trees in all four species.

**Figure 2.**
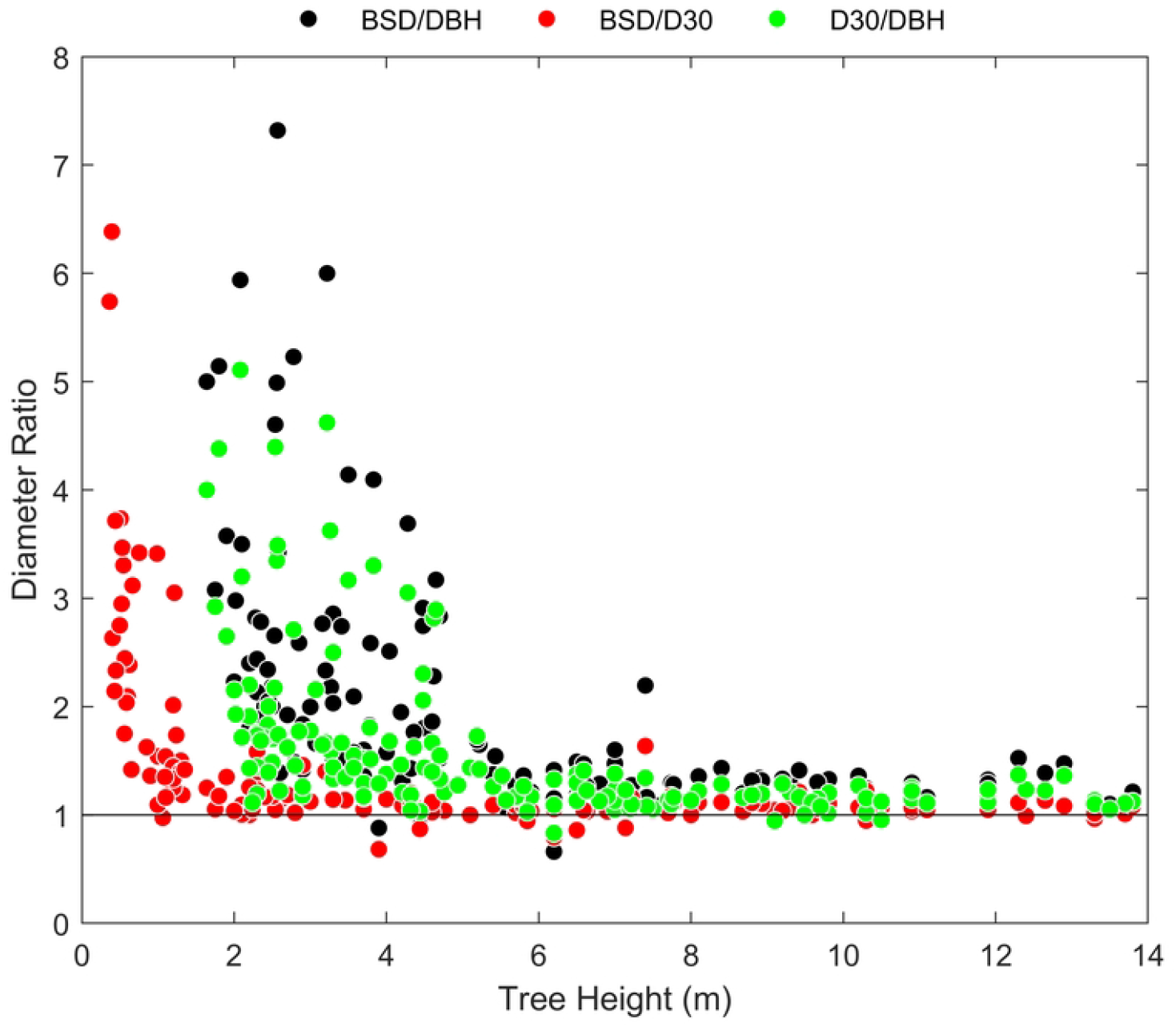
Diameter ratios as a function of tree height. As tree height increases, all three diameter ratios appear to approach a value just above 1, asymptotically. Note that both distributions involving DBH are constrained to begin above a minimum of 1.3 m (DBH height).

**Figure 3.**
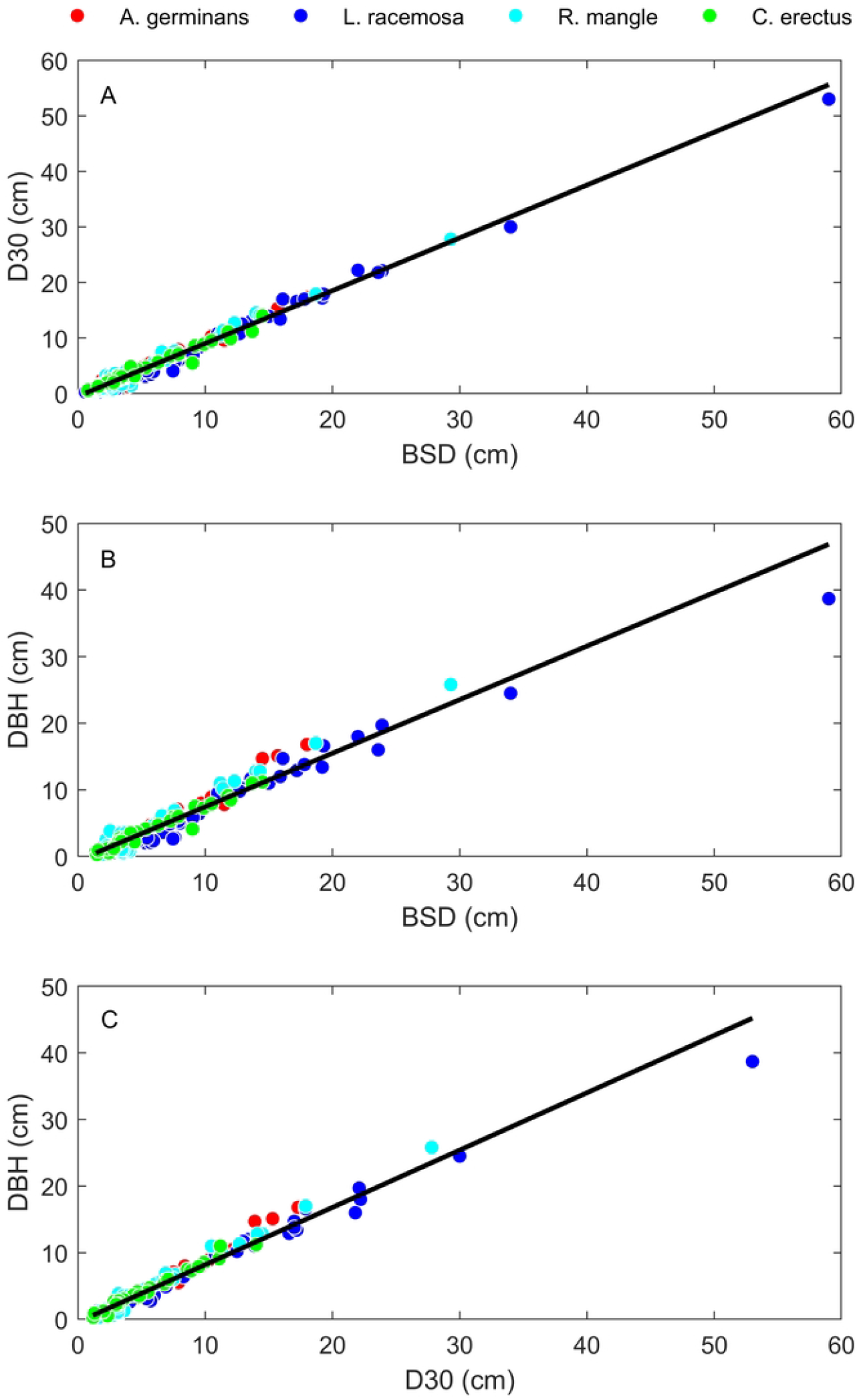
Allometric relationships between the three different diameter measurements. Pairwise bivariate relationships between D30 and BSD (Panel A), DBH and BSD (Panel B), and DBH and D30 (Panel C). Black lines represent interspecific SMA regression lines. Both interspecific and intraspecific regression data are given in Table 3.

Looking further at Table 2, D30 had the lowest RMSE in 17/24 cases, DBH had the lowest RMSE in 4/24 cases, and BSD the lowest in 3/24 cases.

**Table 3.** SMA allometric regression data. Regression statistics for both interspecific and intraspecific bivariate relationships between BSD, D30, and DBH. Note that the slopes for *A. germinans* and *R. mangle* are not statistically different than isometry in all three cases, while those for *C. erectus* and *L. racemosa* are statistically different.

| SMA results |  |  |  |  |  |  |  |  |  |  |
| --- | --- | --- | --- | --- | --- | --- | --- | --- | --- | --- |
| Y variable | X variable | Group | n | R2 | Slope | LowCI | UppCI | Interc | LowCI | UppCI |
| Mean D30 (cm) | Mean BSD (cm) | Interspecific | 238 | 0.986 | 0.9507 | 0.9364 | 0.9652 | -0.4893 | -0.6082 | -0.3704 |
| Mean DBH (cm) | Mean BSD (cm) | Interspecific | 151 | 0.948 | 0.8045 | 0.7753 | 0.8348 | -0.6001 | -0.9057 | -0.2946 |
| Mean DBH (cm) | Mean D30 (cm) | Interspecific | 153 | 0.971 | 0.8607 | 0.8374 | 0.8847 | -0.4177 | -0.6395 | -0.1959 |
| Mean D30 (cm) | Mean BSD (cm) | *A. germinans* | 59 | 0.985 | **0.9906** | 0.9585 | 1.0239 | -0.4751 | -0.6677 | -0.2824 |
| Mean D30 (cm) | Mean BSD (cm) | *C. erectus* | 32 | 0.97 | 0.9036 | 0.847 | 0.9639 | -0.1427 | -0.5348 | 0.2494 |
| Mean D30 (cm) | Mean BSD (cm) | *L. racemosa* | 58 | 0.992 | 0.9356 | 0.9129 | 0.9588 | -0.5893 | -0.8876 | -0.2909 |
| Mean D30 (cm) | Mean BSD (cm) | *R. mangle* | 89 | 0.978 | **1.0269** | 0.9947 | 1.0601 | -0.7851 | -0.9676 | -0.6026 |
| Mean DBH (cm) | Mean BSD (cm) | *A. germinans* | 40 | 0.973 | **1.0012** | 0.9487 | 1.0565 | -1.4401 | -1.8217 | -1.0585 |
| Mean DBH (cm) | Mean BSD (cm) | *C. erectus* | 27 | 0.957 | 0.8176 | 0.7507 | 0.8905 | -0.647 | -1.1558 | -0.1382 |
| Mean DBH (cm) | Mean BSD (cm) | *L. racemosa* | 44 | 0.965 | 0.7509 | 0.7087 | 0.7957 | -0.7035 | -1.353 | -0.0539 |
| Mean DBH (cm) | Mean BSD (cm) | *R. mangle* | 40 | 0.965 | **0.9754** | 0.9171 | 1.0374 | -1.1731 | -1.6515 | -0.6946 |
| Mean DBH (cm) | Mean D30 (cm) | *A. germinans* | 40 | 0.982 | **1.0295** | 0.9851 | 1.076 | -1.1484 | -1.4504 | -0.8463 |
| Mean DBH (cm) | Mean D30 (cm) | *C. erectus* | 27 | 0.979 | 0.9073 | 0.8545 | 0.9633 | -0.5382 | -0.889 | -0.1875 |
| Mean DBH (cm) | Mean D30 (cm) | *L. racemosa* | 45 | 0.982 | 0.8035 | 0.7709 | 0.8374 | -0.2613 | -0.7072 | 0.1845 |
| Mean DBH (cm) | Mean D30 (cm) | *R. mangle* | 41 | 0.981 | **0.9905** | 0.9474 | 1.0355 | -0.9626 | -1.2965 | -0.6287 |

As seen in Table 3, all interspecific slopes were less than isometry. However, half of the intraspecific slopes were not statistically different from the expectation for isometry, with slope 95% confidence intervals that include 1.

## Discussion

Knowledge of the biomass contained in trees at individual, community, landscape, and global levels has been a central focus of ecological research for decades. Model-derived estimates of above ground biomass inform our understanding of community and ecosystem dynamics and guide management decisions. Empirically derived allometric relationships have been a cornerstone of such modelling approaches for over a century [1, 27]

The use of DBH as a biomass predictor has been central to such approaches and is frequently the most informative variable used in predicting tree biomass, when compared to other tree measurements such as tree height or canopy spread [1]. However, there are limitations in using DBH for smaller, shrubbier trees. Foremost among these is the fact that trees in many forests are simply too small to have DBH measured and are typically ignored in censuses or subsampled in small areas. This is despite the fact that small trees can contribute substantially to biomass estimates, particularly in disturbed areas or forest types containing smaller tree species such as mangroves. Further, many trees branch extensively below the DBH measurement point, thus DBH for those trees captures only some of the hydraulic and biomechanical investment required for distal tree branches and is a weaker correlate of total aboveground biomass.

To that end, we compared the predictive power of three different measurements of the diameter of mangrove trees: BSD, D30 and DBH. Looking at the amount of variance explained by the different regression functions (*R*^*2*^), those based on D30 explain more variability than BSD or DBH in ∼71% of cases (Table 2). Moreover, the *R*^*2*^ for both interspecific regressions (within each site and across sites), and the mean *R*^*2*^ for intraspecific regressions are higher for D30 (Table 1). *R*^*2*^ values alone would suggest that D30 is a more adequate variable to measure for biomass estimation.

However, the results using RMSE are somewhat more equivocal. D30 does have the lowest RMSE in ∼71% of the cases, compared to ∼17% for DBH and 12.5% for BSD (Table 2). The percent error relative to the mean mass for the three variables is lower for D30 (10.98%) compared to that for DBH (11.35%) and BSD (15.41% ; Table 2), but only marginally.

Collectively, these results suggest that D30 has a better overall performance than DBH, but not by a lot, and in some cases, at the intraspecific level, DBH has lower RMSE and higher *R*^*2*^ values. We have used *R*^*2*^ and RMSE as complimentary statistics. *R*^*2*^ is informative about the proportion of the variance explained by the regression model whereas RMSE represents the average magnitude of error between the actual and predicted values. Although the two metrics tend to be correlated, it is possible to have a regression model with a high *R*^*2*^ value that also has large RMSE values. This is because *R*^*2*^ is a relative measure of the proportion of total variance, while RMSE is a measure of the absolute difference between the model and the data.

As seen in Fig. 1, the regression functions for all four species are divergent at small tree sizes, but start to converge as trees become larger, with smaller relative differences between diameter measurements (BSD, D30, DBH). This is also reflected in Fig. 2: the diameter ratio values trend larger at small tree sizes, but appear to converge asymptotically to values just above 1 in larger trees. This indicates that the magnitude of the differences in these measurements decreases as trees become large.

All interspecific SMA regression lines had slopes <1 (Table 3). However, for two of the four species (*A. germinans* and *R. mangle*), all three bivariate relationships had intraspecific regression slopes that did not differ from isometry. This suggests the species may differ in shape changes during tree growth. Subsequent inquiries utilizing more comprehensive shape analyses such as with photographs of tree trunks, or terrestrial laser scanning, may provide more insight into the extent to which different mangrove species change shape systematically as they grow, and how this might influence efforts to find reliable proxies for whole plant biomass.

In light of these results we recommend that future studies collecting allometric data for these and other mangrove species consider incorporating multiple measurements of bole diameter so that more data can inform the issue. Since most of the trees in our data compilation are toward the smaller end of the size spectrum, it would be extremely informative if future studies collected these measurements from areas where these species attain their maximum size. At the very least, censuses using D30 will capture more trees due to the lower size threshold. Adding more small trees will help better resolve their contribution to total mass, thus potentially leading to more informed size thresholds for future sampling efforts.

## Acknowledgments

We would like to thank Aerostar Holdings LLC for granting access to the site and for assisting with security clearances and other logistics. Jaime Pabón (Aerostar) provided logistical support and Marisleissis Orona (Aerostar) provided daily escort, supervision of onsite activities, and helped with data acquisition and recording. Walter Soler (Ambienta) assisted with logistical support including the hiring of our chainsaw operator, Alejandro Rodríguez-Rojas (Econet). Ixia Avilés-Vázquez and Luis Ortiz-López (U.S. Forest Service, Forest Inventory and Analysis) also helped with data collection. Funding for SJ Van Bloem provided in part by McIntire Stennis Grant SC-1700590 and Technical Contribution of the Clemson University Experimental Station. Boricorps and the Jobos Bay National Estuarine Research Reserve in Puerto Rico provided field and lab support. Primary funding was provided by U.S. Forest Service Agreement (22-CR-11330145-047). This manuscript was prepared in part by U.S. government employees, therefore it is considered in the public domain and not subject to copyright.

## Supporting Information

**Table S1**. Raw tree measurements and predicted mass. Columns in order contain the dataset, species, mean BSD, D30 and DBH measures, tree height, above ground dry mass (kg), and mean canopy diameter (m). Columns J-U contain the predicted mass based on allometric equations given in Table 1, using interspecific combined data (columns J-L), intraspecific combined data (columns M-O), interspecific site data (columns P-R), and intraspecific site data (Columns S-U).

**Table S2**. OLS regression results for a subset of trees that had all three measurements (BSD, D30 and DBH). The first and second columns contain the Log10 transformed Y and X variables, respectively, followed by the dataset, group, sample size (n), R2 values, slope with lower and upper 95% confidence intervals, and intercept with lower and upper 95% confidence intervals. Thick borders are included to help the reader identify data analyzed at the same level.

